# Control and the Analysis of Cancer Growth Models

**DOI:** 10.1101/244301

**Authors:** Allen Tannenbaum, Tryphon Georgiou, Joseph Deasy, Larry Norton

## Abstract

In this note, we analyze two cancer dynamical models from a system-theoretic point of view. The first model is based upon stochastic controlled versions of the classical Lotka-Volterra equations. Here we consider from a controls point of view the utility of employing ultrahigh dose flashes in radiotherapy. The second is based on work of Norton-Simon-Massagué growth model that takes into account the heterogeneity of a tumor cell population. We indicate an optimal strategy based on linear quadratic control applied to a linear transformed model.

## I. Introduction

In this note, we analyze certain models of cancer growth from a control-theoretic point of view. These include the Lotka-Volterra and Norton-Simon-Massagué models [5], [4], [6], [8], [9], [7]. We are interested in formulating applicable control techniques, so as to formulate more effective therapeutic strategies. In particular, we are interested in exploring from a controls perspective the idea of using ultrahigh dose flashes in radiotherapy [4]. To this end, we first study stochastic controlled versions of the classical Lotka-Volterra equations.

We next turn to the Norton-Simon-Massagué model [6], [8], [9]. This particular approach takes into account the heterogeneity of a tumor cell population following a Gompertzian-type growth curve. A key implication is that therapy should be given at reduced intervals to maximize the probability of tumor eradication and to minimize the chances of tumor regrowth. We will study this more closely via a controlled version of the Norton-Massaguè equation [7].

We will only sketch the basic mathematical theory of the Lotka-Volterra and generalized Gompertzian models used in order to justify our use of certain control laws. A good reference for a more complete mathematical treatment may be found in the text [10] that also has an extensive set of references.

## II. Lotka-Volterra and radiotherapy

We study the following controlled stochastic version for a *competitive Lotka-Volterra* system of equations [10], in order to model the competition between healthy tissue (and/or immune system) on the one hand and cancer cells on the other:

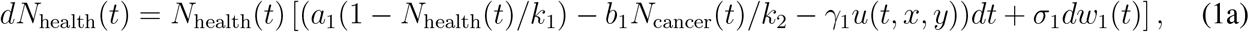

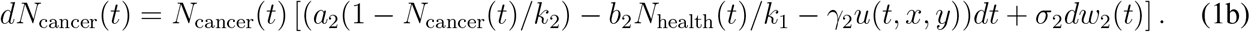

Here, *a_i_* > 0, *k_i_* > 0, *b_i_* ≥ 0, and *γ_i_* ≥ 0 for *i* ∈ {1, 2}, while *u*(*x*, *y*, *t*) represents intensity of radiation which constitutes the control variable. In this model, *N*_health_ represents the number of healthy cells (or the potency of the immune system) and *N*_cancer_ the number of cancer cells. Further

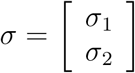

is a 2 × 2 symmetric positive definite matrix (i.e., *σ_i_* are 1 × 2 row vectors), and

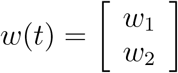

is a 2-dimensional Brownian motion (Wiener process) that models randomness. The parameters *γ_i_* represent the susceptibility of the respective populations to radiation, the parameters *a_i_*, *b_i_* represent the rates of growth and interaction between the two populations respectively, while *k_i_*, for *i* ∈ {1, 2}, represents the saturation value for the respective population. Typically *a*_2_ >> *a*_1_ and *k*_2_ >> *k*_1_. Without loss of generality we may assume that *γ*_2_ = 1 and therefore redefine *γ*_1_ =: *γ* to simplify notation.

We normalize the above populations by setting

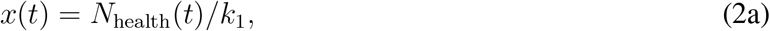

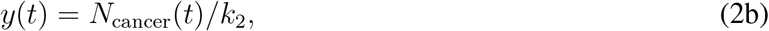

and thereby we re-write the system dynamics as follows:

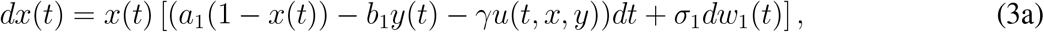

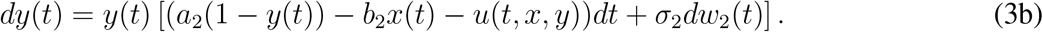

The problem we consider is to choose a control strategy (i.e., shape and duration of the radiation intensity *u*(*t*, *x*, *y*)) so as to reduce the tumor from a given starting point

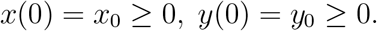

We will consider both open loop as well as closed loop control. The latter assumes an online estimate of the effectiveness of radiation.

One particular question that we set to answer is the question of whether, given the total amount of radiation, ***it is preferable to deliver the treatment over a wide window of time or a short one with high intensity***.

Interestingly, depending on model parameters, there are cases where cancer always wins out unless ameliorated by control. Those cases require periodic treatment in perpetuity. For other model parameters, there are typically two regions for (*x*, *y*). In the first region, where concentration of cancer cells is significant, cancer wins out. In the second, the relative proportion of cancer cells is insignificant and the immune system is seen as capable of eliminating all cancerous cells. Accordingly,

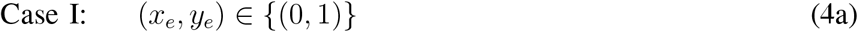

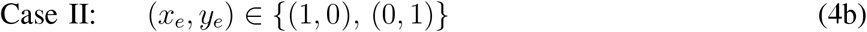

are the corresponding points of equilibrium. The corresponding values for *N*_health_, *N*_cancer_ are *k*_1_, *k*_2_, respectively.

### A. Analysis of system

We first consider and analyze the deterministic and autonomous system:

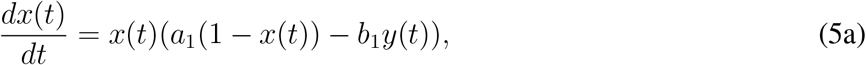

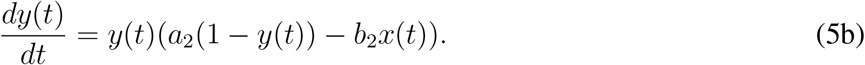

In general, there are three possible points (*x_e_*, *y_e_*) of equilibrium,

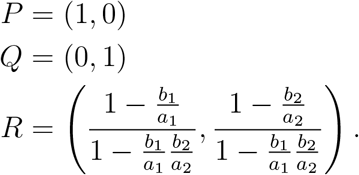

Linearization of the dynamics about these points of equilibrium gives

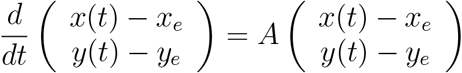

where *A* is a 2 × 2 matrix taking values

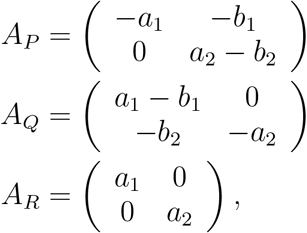

respectively. Thus, *P* and *Q* are stable equilibria provided *a*_1_ < *b*_1_ and *a*_2_ < *b*_2_, respectively, whereas *R* is always unstable.

Thus, without control, in this deterministic case, cancer cells will always win the competition when *a*_1_ < *b*_1_ < 0, and *a*_2_ > *b*_2_. In this case, *N*_health_(*t*) → 0 and *N*_cancer_(*t*) → *k*_2_ as *t* → ∞, or equivalently, *x*(*t*) → 0 and *y*(*t*) → 1.

### B. Case I: cancer always wins without control

In the simulation of an academic example below, we chose *k*_1_ = *k*_2_ = 1, *a*_1_ = 0.01, *a*_2_ =0.2, *b*_1_ = 0.02, *b*_2_ = 0.1, *x*_0_ = *y*_0_ = .5 and control input 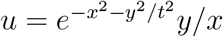. This gives an impulsive nature to the control.

The control contains a burst and then continues on at some level, otherwise cancer will re-appear. See Figures 1 and 2.

**Fig. 1.**
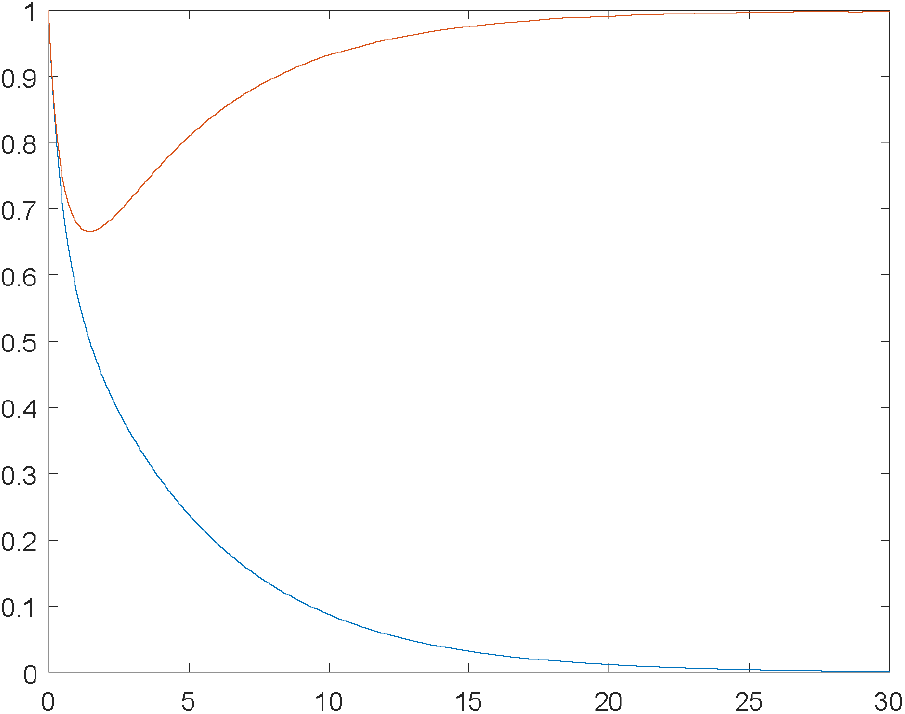
Cancer/Normal Cell Concentrations: Time series of Lokta-Volterra system with no control applied. Red is cancer and blue is healthy.

**Fig. 2.**
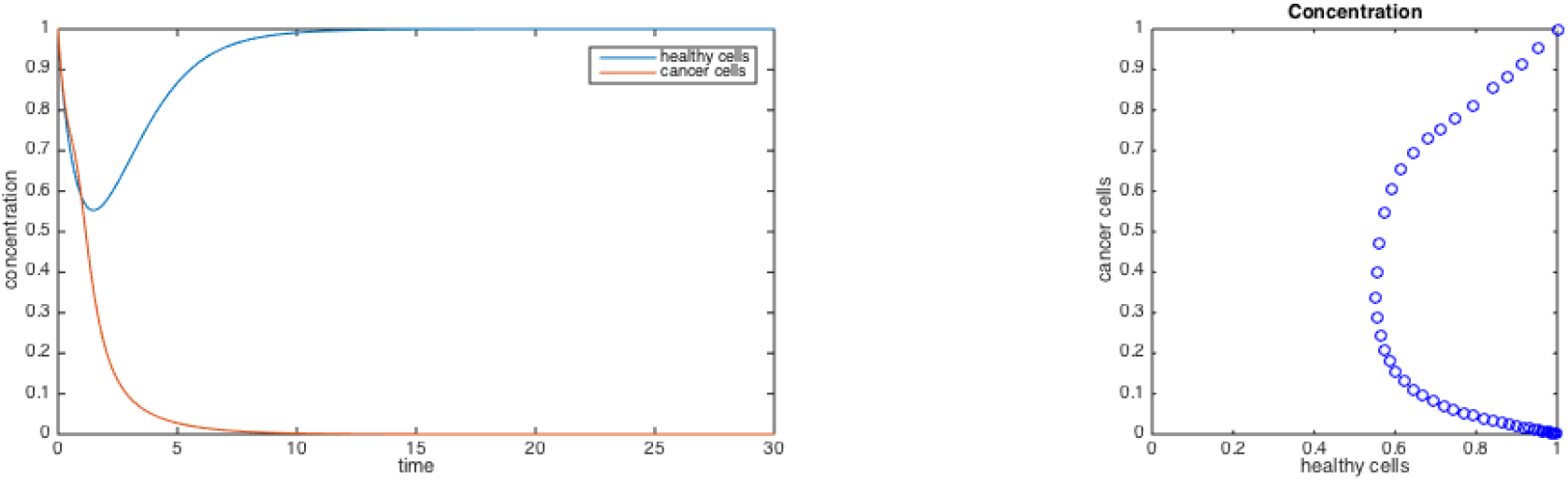
Cancer/Normal Cell Concentrations: i) time series, ii) phase plot of Lokta-Volterra system with impulsive control applied. Red is cancer and blue is healthy.

## III. Case II: if tumor sufficiently shrinks, it is eliminated by the body

In this case, we assume that both *P* and *Q* are attractive points of equilibrium and, therefore, proper radiation treatment has a chance to lead the dynamics towards *P* which represents cure. However, the region in phase space (*x*, *y*) where *P* is an attractive equilibrium and thereby the immune system is capable of eliminating cancer, could be very small. The task then is to steer the system into this region with suitable radiation treatment.

We demonstrate that if a certain amount of radiation is allowable to be delivered (i.e., intensity × duration = fixed), then it is advantageous to deliver all the radiation in the shortest possible amount of time. Below, we present a numerical example where we assume *a*_1_ = 1, *a*_2_ = 1, *b*_1_ = 2, *b*_2_ = 1.1, *γ* = 0.2, *u* constant over an interval *T* such that *u* × *T* = 3. We show phase plots marking the path of the concentrations (*x*(*t*)*,y*(*t*)) starting from (0.7333, 0.3833) for a range of values for the intensity *u* and duration *T* as follows

| duration | intensity |
| --- | --- |
| 15 | 0.2 |
| 10 | 0.3 |
| 7.5 | 0.4 |
| 6 | 0.5 |
| 5 | 0.6 |
| 4.2857 | 0.7 |
| 3.75 | 0.8 |
| 3 | 1 |

We observe that short duration and higher intensity treatment is preferential since it is more effective in steering the state into the region of attraction of the point of equilibrium *P* representing cure. See Figures 3 and 4.

**Fig. 3.**
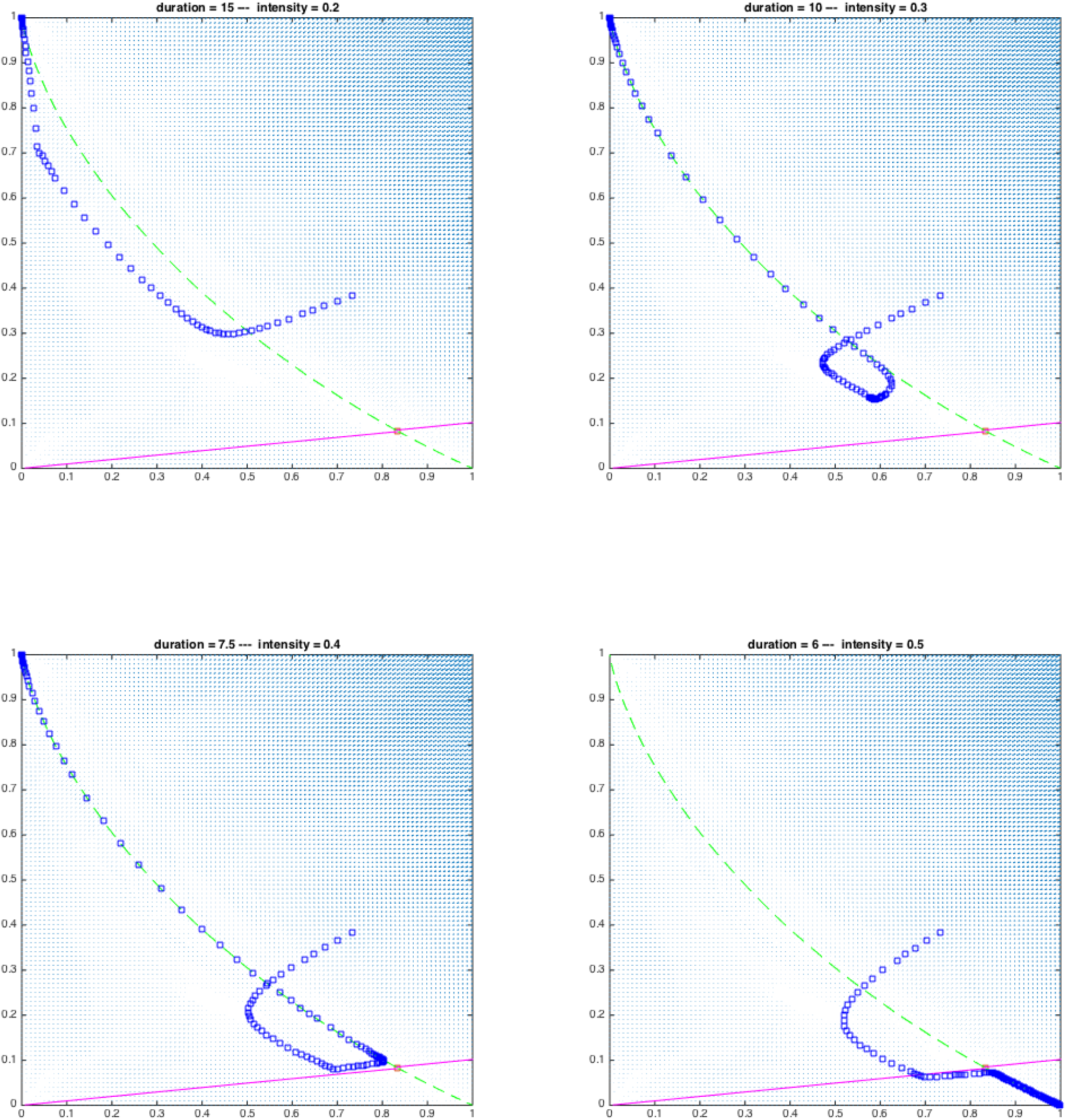
Cancel/Normal Cell phase diagram (longer duration/weak intensity)r: In these simulations, we start at the unstable equilibrium point. Given its sensitivity, we plot both the forward (green dashed) and time-reversed (magenta) trajectories of the Lotka-Volterra system so they intersect at the unstable equilibrium. Note that for very small changes in the initial condition, the forward trajectory can either go the cancerous equilibrium point (0, 1) or the healthy point in which the cancer is eradicated (1, 0). Here we added a control of relatively weak intensity and longer duration. Note that the system does not converge to the healthy point (1, 0) in the first three plots (blue curves). Only when we increase the intensity is the system driven the system to (1, 0) in the last plot. In Figure 4 we continue this analysis, with short duration inputs of higher intensity.

**Fig. 4.**
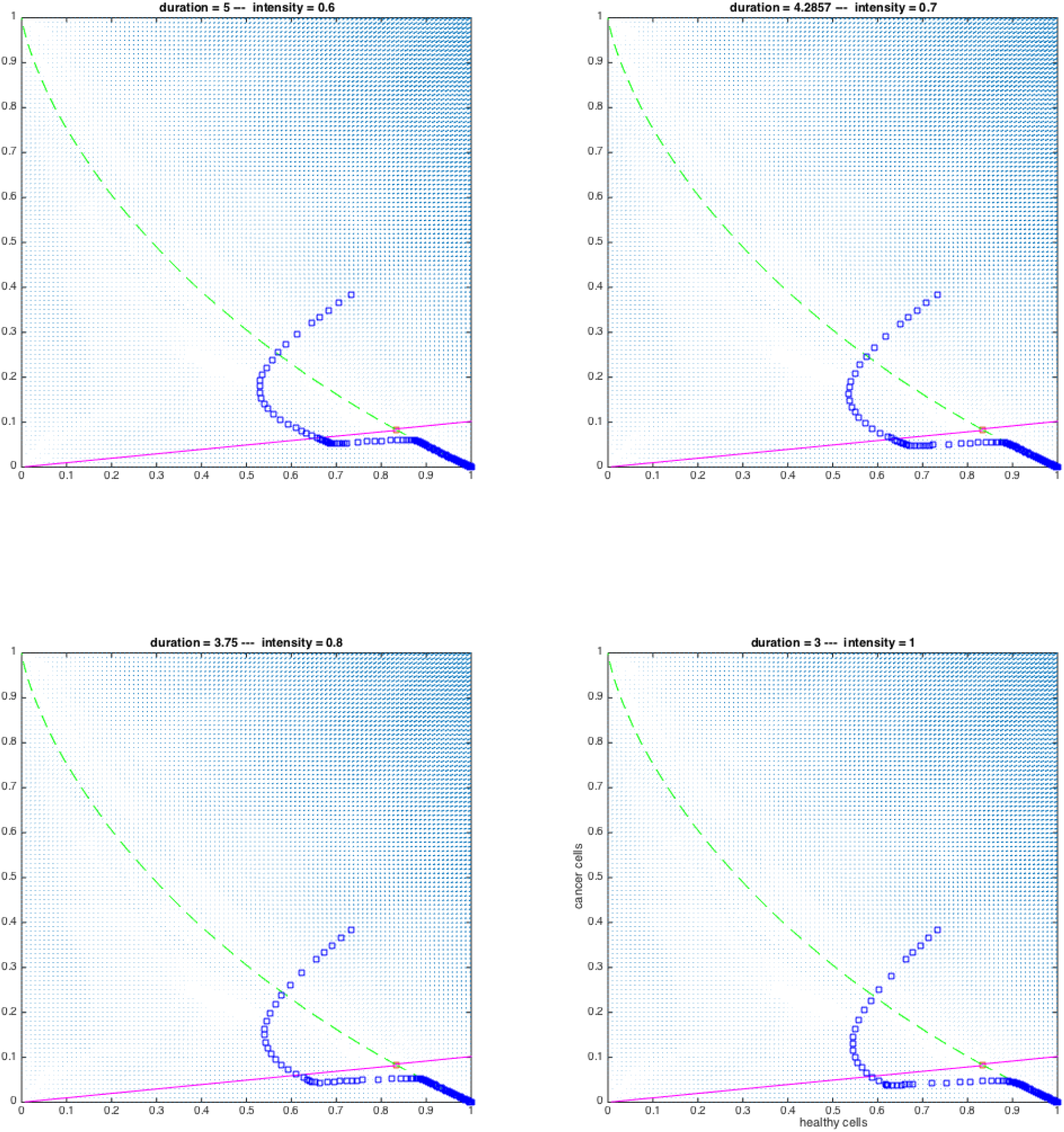
Cancer/Normal Cell phase diagram (short duration/high intensity): In these simulations, we start at the unstable equilibrium point. Given its sensitivity, we plot both the forward (green dashed) and time-reversed (magenta) trajectories of the Lotka-Volterra system so they intersect at the unstable equilibrium. Note that for very small changes in the initial condition, the forward trajectory can either go the cancerous equilibrium point (0, 1) or the healthy point in which the cancer is eradicated (1, 0). Adding control of a high intensity and short duration drives the system to the healthy point (1, 0) (blue curves).

## IV. Norton-Massagué dynamics: system identification and control

A key hypothesis in the Norton-Simon-Massagué models [6], [8], [9], [7], which is supported by experimental evidence, is that small tumors grow faster, but they are also more susceptible to treatment. The accentuated effect of small-size in growth rate is captured by a fractional power <1 in population dynamics, as in

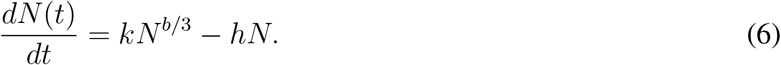

We will refer to this as the Norton-Massagué equation. There are several simple observations to make. First, small *b* gives a rather high slope near *N* = 0. In this case, *N* will not go to infinity. When *b* > 3 the system will blow up, but for small *N*, linear dissipation dominates.

We propose to add a control aspect to the model. For example, the effect of control *u* (effect of radiation or drugs) on such an evolution could be modeled via

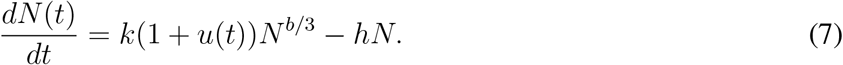

The choice of *u* specifies the control strategy. When optimizing a quadratic cost, the optimal control is typically in the form of state feedback, i.e., a function of *N*(*t*).

We point out that the Norton-Massagué equation may be solved in closed form, namely,

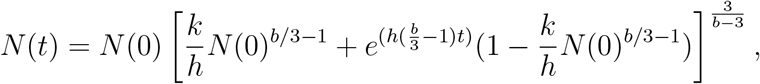

as may be checked by direct differentiation. Indeed, the Norton-Massagué equation is a special case of a Bernoulli equation [2], i.e., an equation of the form

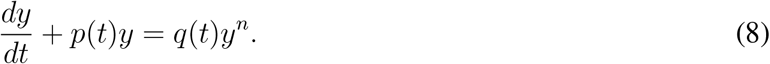

In the Norton-Massagué case, *y*(*t*) = *N*(*t*), *p*(*t*) = *h*, *q*(*t*) = *k*, and *n* = *b*/3.

Specifically, via a simple transformation equation (8) may be converted into a linear equation. We set *y*′ = *dy*/*dt*. Now divide both sides of (8) by *y^n^* to get

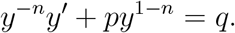

Set *v* := *y*^1−^*^n^*, so that *v*′ = (1 − *n*)*y*^−^*^n^y*′. Therefore, (8) becomes

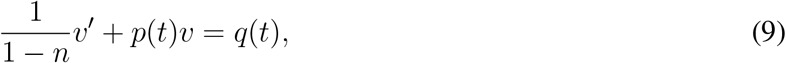

which is linear. This linear model motivates our choice of the control strategy for the Norton-Massagué equation.

### A. Optimal Control

In this section, we apply a linear quadratic regulator performance index [1] to the Norton-Massagué equation. Set *n*(*t*) := *N*(*t*)*^c^*, *c* := 1 − *b*/3.. Then *n*(*t*) satisfies the equation

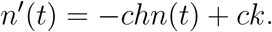

We modify this equation by adding a control *u*(*t*)

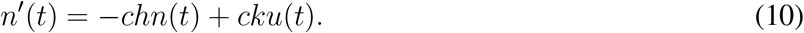

A suitable performance index to be minimized over choice of control-input in equation (10) is:

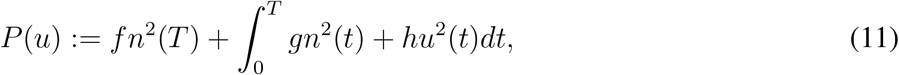

where the time *T* is the therapeutic treatment horizon, and *f*, *g*, *h* > 0 are parameters to be chosen. Thus, we seek to minimize *P*(*u*) over all piecewise continuous functions *u* : [0,*T*] → [0,*u_max_*] subject to the dynamical constraint given by equation (10). Here *g* weights the number of cancer cells remaining at the end of treatment, and *h* scales the running cost that measures the tumor volume. It is well-known that the optimal control has the form *u*(*t*)= −*α*(*t*)*n*(*t*) where *α* may be found via a Riccati equation [1]. When transformed back to the original Norton-Massagué model, this leads to a control law of the form *u*(*t*)= −*α*(*t*)*N*(*t*)*^c^*.

### B. State Feedback

For the controlled Norton-Massagué equation (7), and for time-varying *u*(*t*), we get the following explicit solution:

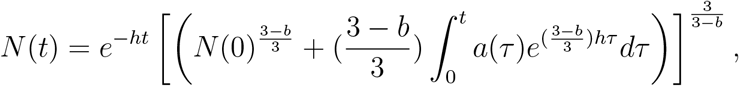

where *a*(*t*) := *k*(1 + *u*(*t*)).

### C. Numerical Example

We illustrate the use of state feedback in which the state of the system is used to close the feedback loop by desplaying responses of the (controlled and uncontrolled) system in Figure 5 below. In a realistic scenario, one would need an estimator as well since the signals would be corrupted by noise. Here we show the following: we start with a small tumor (*N*(0) = 1) and take *b* = 2.5 in the Norton-Massague equation. As expected, we get an initial sharp increase, and then much slower growth. We then assume that the exponent *b* becomes 3.1, and as expected without control, one gets rapid exponential growth. In the final plot, we show what what happens if we add feedback when *b* = 3.1. The tumor growth is controlled. Note that we use the approximate steady state of the Norton-Massague equation for *b* = 2.5 as the input to the controlled and uncontrolled models.

**Fig. 5.**
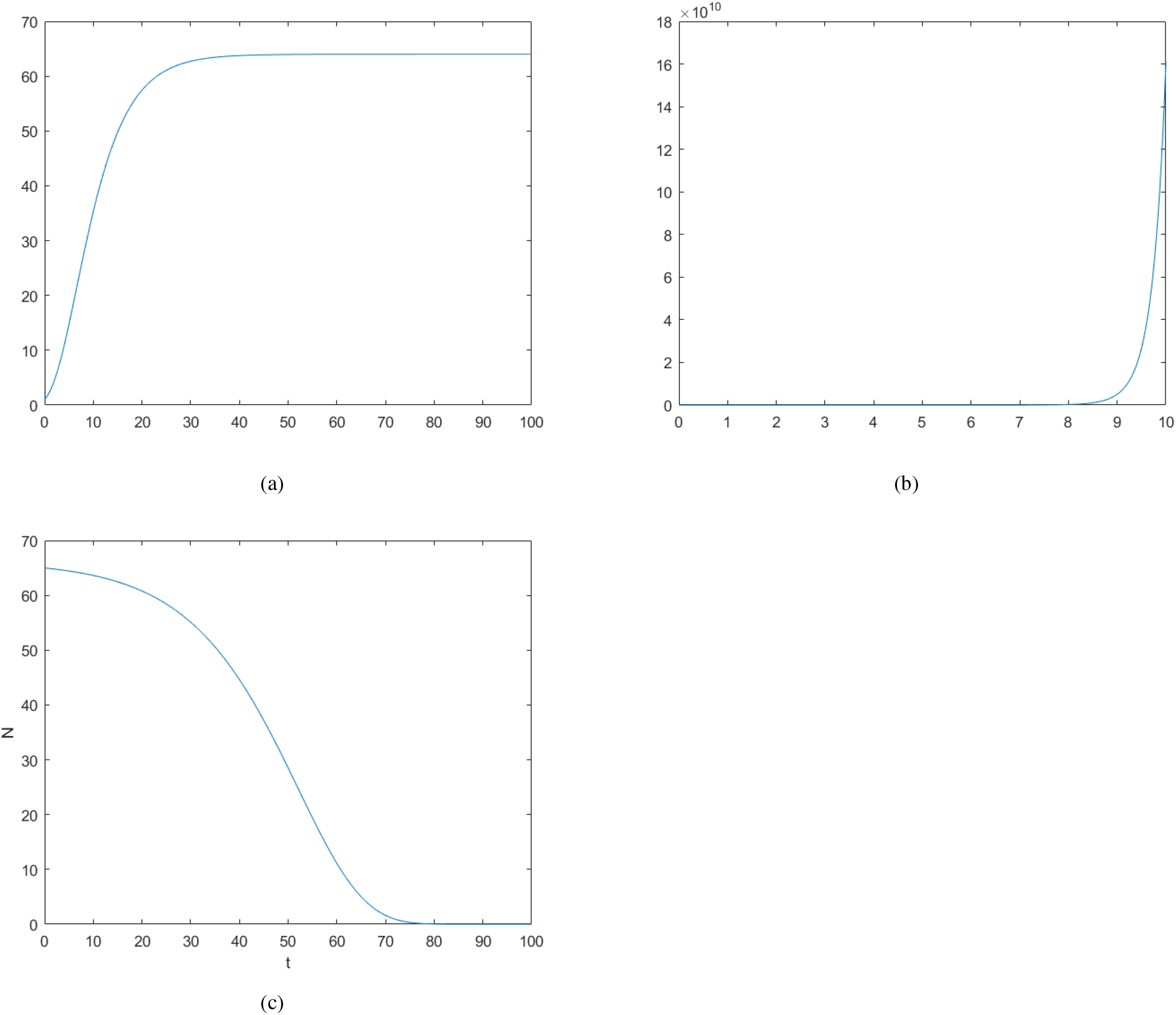
Top Row: Growth via the Norton-Massagué equation with *b* = 2.5 and *b* = 3.1 without control; bottom figure: *b* = 3.1 with state feedback

## V. Conclusions

The results of this note are of course very preliminary. The note sets down some very basic ideas from systems and control that we would like to exploit for developing more effective cancer therapies. In particular, we plan to use ideas from system identification backed up by laboratory experiments, in order to discover and tune more precisely the relevant parameters in our dynamical evolution equations. We have not included noise in our simulations so far. We plan to develop suitable noise models that will drive the stochastic equations. At Sloan Kettering, we have access to a great deal of CT lung tumor data before and after radiotherapy, that could be very useful in this regard.

Finally, for the Norton-Massagué model we have only explored the controls that are optimal with respect to a “linear-quadratic” performance index. We plan to employ more sophisticated ideas from robust control that would in addition ensure small sensitivity to parameter uncertainly [3].

## Acknowledgments

This work was supported by the Breast Cancer Research Foundation.

